# Identification and *in silico* analysis of the origin recognition complex in the human fungal pathogen *Candida albicans*

**DOI:** 10.1101/430892

**Authors:** Sreedevi Padmanabhan, Kaustuv Sanyal, Dharani Dhar Dubey

**Affiliations:** Molecular Biology Laboratory, Department of Biotechnology, Veer Bahadur Singh Purvanchal University, Jaunpur 222 003, Uttar Pradesh, India; Molecular Mycology Laboratory, Molecular Biology and Genetics Unit, Jawaharlal Nehru Centre for Advanced Scientific Research, Jakkur, Bangalore 560 064, India.

## Abstract

DNA replication in eukaryotes is initiated by the orchestrated assembly and association of initiator proteins (heterohexameric Origin Recognition Complex, ORC) on the replication origins. These functionally conserved proteins play significant roles in diverse cellular processes besides their central role in ignition of DNA replication at origins. While *Candida albicans*, a major human fungal pathogen, is an ascomycetous, asexual, diploid budding yeast but it is significantly diverged from a much better studied model organism *Saccharomyces cerevisiae*. The components of the DNA replication machinery in *C. albicans* remain largely uncharacterized. Identification of factors required for DNA replication is essential for understanding the evolution of the DNA replication machinery. We identified the putative ORC homologs in *C. albicans* and determined their relatedness with those of other eukaryotes including several yeast species. Our extensive *in silico* studies demonstrate that the domain architecture of CaORC proteins share similarities with the ORC proteins of *S. cerevisiae*. We dissect the domain organization of ORC (trans-acting factors) proteins that seem to associate with DNA replication origins in *C. albicans*. We present a model of the 3D structure of CaORC4 to gain further insights of this protein’s function.

## Introduction

DNA replication in eukaryotes is initiated by the orchestrated assembly and association of initiator proteins on the replication origins. The hunt for initiator proteins in higher eukaryotes picked up pace after the discovery of the Origin Recognition Complex (ORC) comprising of six protein subunits of the ORC1-6 complex in budding yeast^1^. Extensive studies in other organisms showed that the initiator ORC proteins are functionally conserved in all eukaryotes and the association of ORC proteins with DNA replication origins is critical for initiation of DNA replication, a fundamental process of life. The replicators, occupied by ORC proteins, fire asynchronously in S phase. Replicators in different organisms have widely variable DNA sequence requirements. In some organisms no obvious DNA sequence requirements could be detected. The sequential assembly of the pre-replication complex (pre-RC) proteins on the origins is mediated by ORC. ORC orthologs have been identified in many eukaryotes like *Schizosaccharomyces pombe, Drosophila melanogaster, Xenopus laevis*, and humans. Genetic and biochemical investigations demonstrate the ORC proteins of these organisms to be essential for DNA replication initiation^2^. Although the replication origins in higher eukaryotes do not share a consensus sequence as in bacteria or budding yeast, the proteins that are recruited to origins in most metazoans are similar to those in bacteria and yeast suggesting replication-associated proteins are evolutionarily conserved^3–4^. ORC-mediated ATP hydrolysis is essential for recruiting MCM (Mini Chromosome Maintenance) proteins^5^ which subsequently act as helicases and unwind DNA double helix to facilitate initiation of DNA replication.

Besides playing a central role in DNA replication initiation at discrete origin sites, ORC proteins are also involved in a variety of cellular processes like heterochromatin formation, transcriptional regulation, S-phase checkpoint regulation, mitotic chromosome assembly, sister chromatid cohesion, cytokinesis, ribosome biogenesis and tissue specific gene regulation. ORC mutations are seen in various human diseases^6^ such as Meier–Gorlin syndrome^7, 8^, EBV (Epstein–Barr virus)-infected diseases^9^, American trypanosomiasis and African trypanosomiasis^10^.

There has been a wide prevalence of yeast infections over the years with *Candida* species that can cause superficial to fatal systemic infections. These fungal infections can be fatal for immunocompromised individuals where the mortality rate is even higher^11^. Availability of a handful antifungal drugs and frequent isolation of drug-resistant isolates led to complications in disease management and treatment procedures^12^. Hence, to circumvent this malicious infection is to find species-specific new drug targets to develop more effective and safer antifungals. *C. albicans* is one such opportunistic fungal pathogens which is an asexual, diploid, budding yeast^13–14^. Protein complexes involved in the DNA replication of *C. albicans* are not characterized. As DNA replication is a rate limiting step in the propagation of the yeast and not many anti-fungal drugs are available to curb *Candida* infection, in this study we sought to identify and dissect the domain architecture of CaORC proteins with an aim to provide clues on their evolutionary conservation/diversification across various species to explore their potential as species-specific drug targets.

## Results

### Identification of preRC genes in *C. albicans* genome by *in silico* analysis

First, we identified the homologs of the preRC complex in *C. albicans*, determined relatedness of these proteins present in other species, compared the various domains such as the BAH domain, AAA+, AT-hook and Walker motifs in the ORC proteins of a number of species and predicted the structure of ORC4 in *C. albicans*, CaORC4. The CaORC1-6 genes were identified by a BLAST search with *S. cerevisiae* ScORC1-6 as the query sequences against the *C. albicans* genome database (CGD) (http://www.candidagenome.org/cgi-bin/compute/blast-sgd.pl) (Table 1 and Table 2).

**Table 1.**
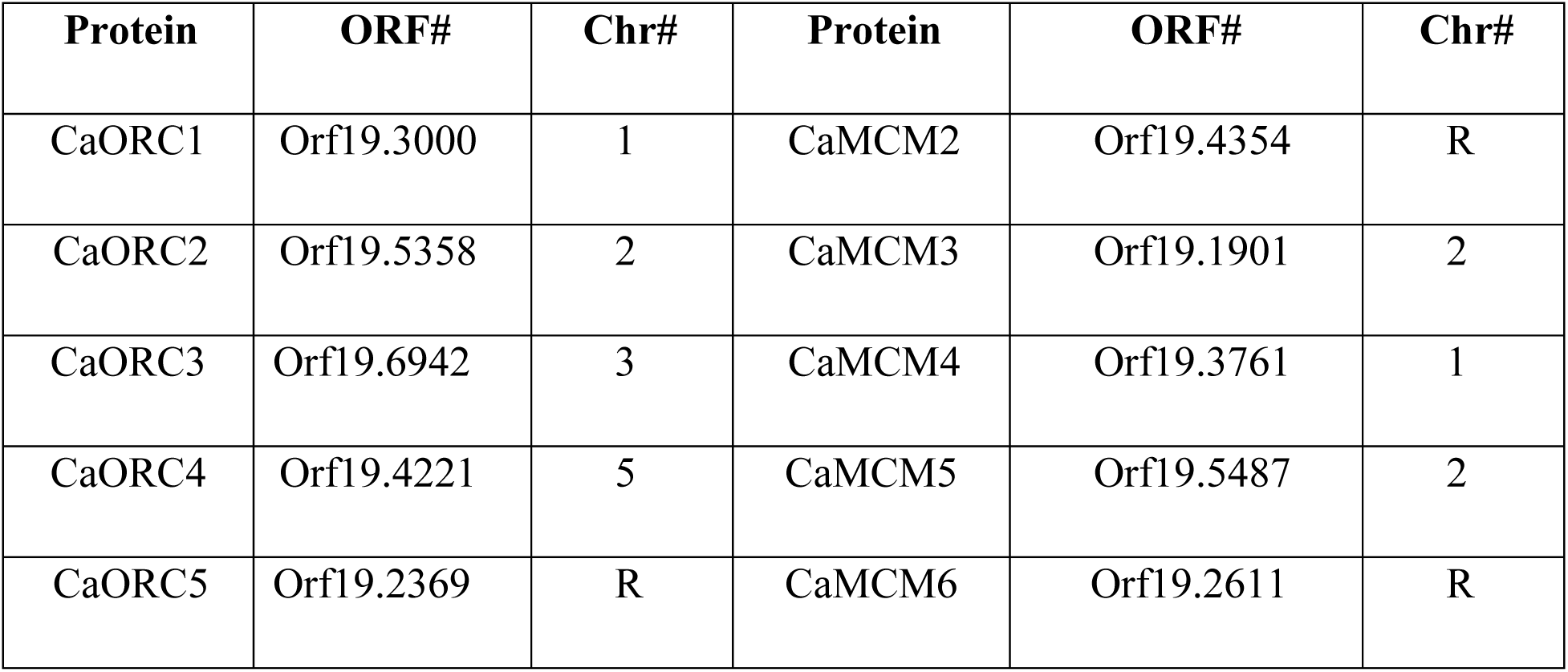
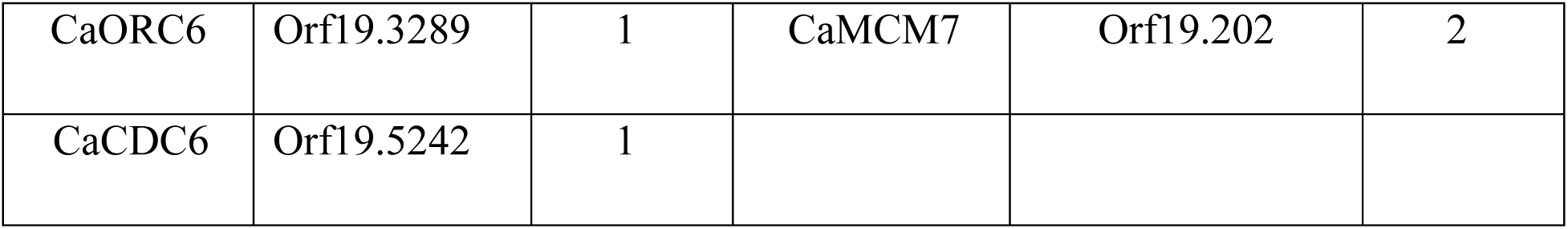
Putative pre-RC proteins coded by the *C. albicans* genome

**Table 2.**
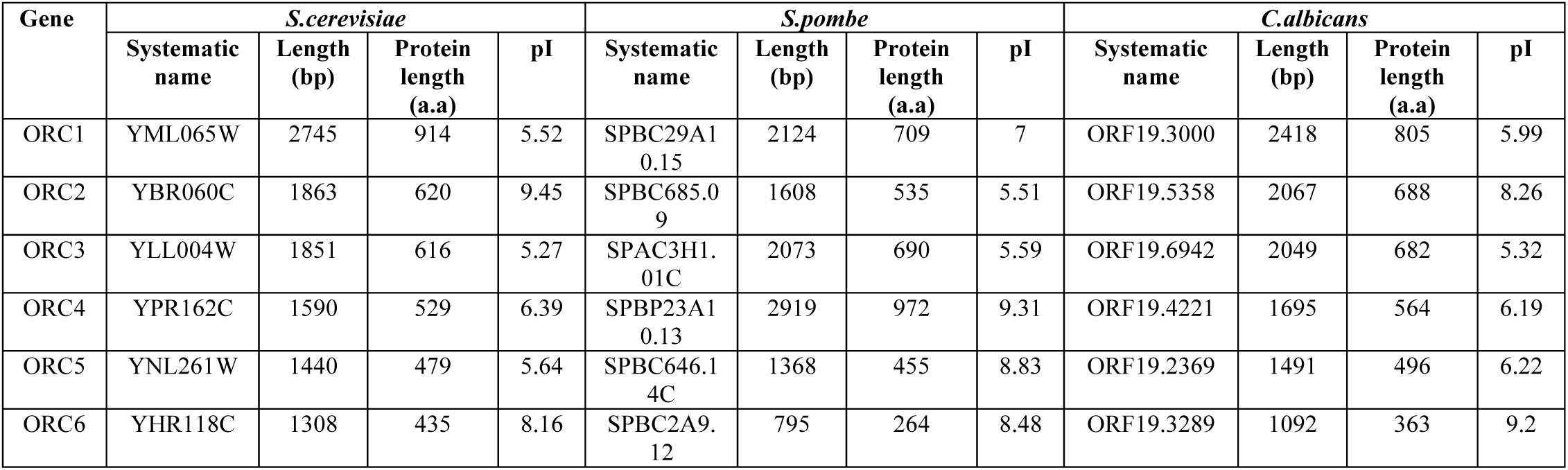
Comparison of putative CaORC sequences with ORC sequences of *S. cerevisiae* and *S. pombe*. The systematic names, ORF and protein length along with isoelectric pH of the ORC1-6 in *S. cerevisiae, S. pombe* and *C. albicans*.

ClustalW2 is a DNA or protein multiple sequence alignment program for multiple sequences^15^. We performed a pair-wise amino acid sequence alignment of the CaORC proteins with those of *S. cerevisiae, S. pombe, Drosophila, Xenopus*, Mouse and humans individually and their respective clustalW scores are tabulated (Table 3). Although, in general, the CaORC proteins show limited sequence similarities with their counterparts in various species, CaORC1, 2 and 6 show maximum similarities to their *S. cerevisiae* counterparts while CaORC3, 4 and 5 appear to be more similar to those of mammals which is evident from the phylogenetic map (Figure 1A).

**Table 3.** Pairwise alignment results of CaORC proteins with other eukaryotes.

| Protein Name | Clustal W scores |  |  |  |  |  | Length of protein (a.a) |  |  |  |  |  |  |
| --- | --- | --- | --- | --- | --- | --- | --- | --- | --- | --- | --- | --- | --- |
|  | *Ca vs Sc* | *Ca vs Sp* | *Ca vs Dm* | *Ca vs Xl* | *Ca vs Mm* | *Ca vs Hs* | *Ca* | *Sc* | *Sp* | *Dm* | *Xl* | *Mm* | *Hs* |
| ORC1 | 26 | 26 | 20 | 20 | 23 | 20 | 805 | 914 | 709 | 924 | 886 | 840 | 861 |
| ORC2 | 25 | 21 | 19 | 16 | 20 | 19 | 688 | 620 | 535 | 618 | 558 | 576 | 577 |
| ORC3 | 18 | 14 | 13 | 14 | 18 | 17 | 682 | 616 | 690 | 721 | 709 | 715 | 712 |
| ORC4 | 25 | 24 | 21 | 23 | 25 | 23 | 564 | 529 | 972 | 459 | 432 | 433 | 436 |
| ORC5 | 22 | 21 | 20 | 17 | 23 | 24 | 496 | 479 | 455 | 460 | 448 | 435 | 435 |
| ORC6 | 17 | 12 | 6 | 15 | 9 | 11 | 363 | 435 | 264 | 257 | 225 | 262 | 252 |

*Sc* – *Saccharomyces cerevisiae*; *Sp* – *Schizosaccharomyces pombe*; *Ca* – *Candida albicans*; *Dm* – *Drosophila melanogaster*; *Xl* – *Xenopus laevis*; *Mm* – *Mus musculus*; *Hs* – *Homo sapiens*

**Figure. 1.**
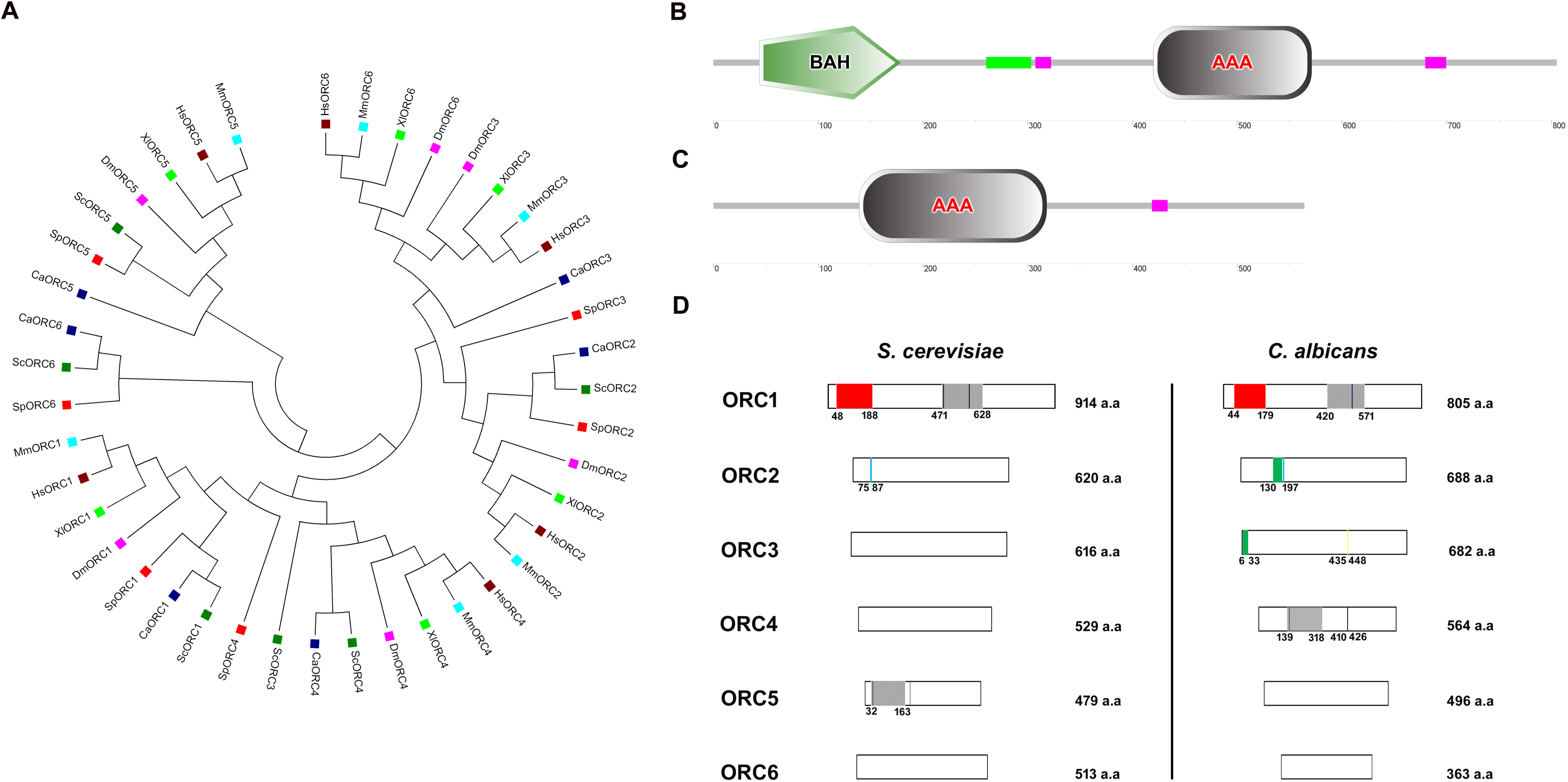
Evolutionary relationship of CaORC proteins with other species and comparative domain architecture of CaORC and ScORC proteins. **(A)** Phylogram of ORC proteins. The tree is drawn to scale, with branch lengths in the same units as those of the evolutionary distances used to infer the phylogenetic tree. The evolutionary distances were computed using the Poisson correction method^77^ and are in the units of the number of amino acid substitutions per site. All positions containing gaps and missing data were eliminated from the dataset (Complete deletion option). There were a total of 116 positions in the final dataset. Phylogenetic analyses were conducted in MEGA4^35^. **(B)** The SMART (Simple Modular Architecture Research Tool) prediction shows the presence of the BAH domain spanning between 44^th^ and 179^th^ amino acids at the N-terminal of CaORC1 and **(C)** The AAA+ domain in CaORC4 protein, the purple box represents the low complexity region (LCR). The LCR may be involved in flexible binding associated with specific functions but also that their positions within a sequence may be important in determining both their binding properties and their biological roles ^78^. **(D)** Comparative domain architecture of ORC proteins in *S. cerevisiae* and *C. albicans*. The red box denotes the BAH domain, the grey box is the AAA+ domain, cyan bar represents the AT-hook motif, black bar represents the Walker motifs, dark blue bar represents the PIP motif, yellow bar represents the MIR motif and the green bar represents the PEST motif.

### BAH domain in ORC proteins

The Expasy PROSITE consists of documentation entries describing protein domains, families and functional sites as well as associated patterns and profiles to identify them. The Expasy PROSITE tool predicts the presence of an evolutionarily conserved BAH_domain spanning the region between 44^th^ and 179^th^ amino acids at the N-terminal of CaORC1 (Figure 1B). The BAH domain is involved in protein-protein interactions and has been found to be important in DNA methylation, replication and transcriptional regulation^16^.

### AAA+ domains in CaORC proteins

<u>A</u>TPases <u>a</u>ssociated with various cellular <u>a</u>ctivities (AAA+) domains^17–18^ are those that are activated by ATP binding and inactivated by ATP hydrolysis^19–23^. The ATPase activity is indispensable for the origin-ORC association and henceforth for the establishment of the pre-initiation complex. Preventing ORC ATP hydrolysis inhibits repeated MCM2-7 loading^5^. CaORC1 and CaORC4, each contains a consensus AAA+ domain (420-571 a.a. in CaORC1; 139-318 a.a. in CaORC4) (Figure 1B, 1C and 1D), which belongs to the AAA+ family that is pivotal to the initiation of eukaryotic DNA replication. There is an amino acid residue Tyr^174^ in human ORC4 (Tyr^232^ in *S. cerevisiae)* that is found between the Walker B motif and sensor I of the AAA+ domain which may be responsible for interacting with a conserved arginine residue on an adjacent helix structure of ORC4^2,6,21,23–26^. This residue is present in CaORC4 (Tyr^273^) too probably doing a similar function.

Although the ScORC1-ORC5 all have AAA+ domains, there is a variation among the subunits with respect to the catalytic core motifs within the Walker A and B motif regions both within and between the species. It is reported with experimental evidence that only the ScORC1 and ScORC5 can bind ATP, of which only ScORC1 has a perfect signature to the Walker B motif. By consensus, it is considered that the ORC1 would be the prime ATPase of all the eukaryotes examined so far^25, 26^. In metazoans too, although the ORC1, ORC4 and ORC5 bind ATP, the Walker A signature is found to have perfect match with ORC4. A similar pattern is observed in CaORC proteins too demonstrating their close homology with the higher eukaryotes. The Walker A motifs in ORC5 seem to be diverged (Table 4).

**Table 4.** Comparison of Walker A and Walker B motifs of CaORC proteins with other species

| Protein name | Organism | Walker A motif<br>(GXXXXGKT/S) | Walker B motif<br>(hhDE) |
| --- | --- | --- | --- |
| **ORC1** | *S. cerevisiae* | GTPGVGKT | LLDE |
|  | *S. pombe* | GTPGTGKT | LMDE |
|  | *C. albicans* | GVPGMGKT | LMDE |
|  | *C.elegans* | GVPGTGKT | LI DE |
|  | *D. melanogaster* | GVPGTGKT | LVDE |
|  | *M. musculus* | GVPGTGKT | LVDE |
|  | *H. sapiens* | GVPGTGKT | LVDE |
| **ORC4** | *S. cerevisiae* | GPRQSYKT | IFDE |
|  | *S. pombe* | GPRGSGKS | VLEE |
|  | *C. albicans* | GPRSSGKT | SDLE |
|  | *C.elegans* | GERNCGRE | LVRD |
|  | *D. melanogaster* | GPRGSGKT | ILEE |
|  | *M. musculus* | GPRGSGKT | ILDE |
|  | *H. sapiens* | GPRGSGKT | ILDE |
| **ORC5** | *S. cerevisiae* | GYSGTGKT |  |
|  | *S. pombe* | GVASTAKT |  |
|  | *C. albicans* | GYKSIGKT |  |
|  | *C.elegans* | GEDGSGRS |  |
|  | *D. melanogaster* | GHSGTGKT |  |
|  | *M. musculus* | GHTASGKT |  |
|  | *H. sapiens* | GHTASGKT |  |

### Walker A and B motifs in CaORC proteins

The motif GXXXXGKT (X, any residue) is a common nucleotide binding fold in the α- and β-subunits of F1-ATPase, myosin and other ATP-requiring enzymes^27^. This motif is present in the shape of a loop around nucleotides and utilizes its highly conserved residues of lysine and threonine to bind to their phosphate oxygen atoms. This consensus sequence of GXXXXGKT(S), with serine substituting threonine in some cases, is more popularly known as the Walker loop or P-loop (phosphate binding loop). The Walker B motif with the consensus sequence hhhhDE (a negatively charged residue followed by a stretch of hydrophobic a.a,,h) is essential for ATP hydrolysis. The Walker motifs are present in CaORC1, CaORC4 and CaORC5 (Walker B is absent in CaORC5) (Table 4). Besides CaORC1, the perfect signature of the Walker motif is found in CaORC4 with a putative Walker A motif (147–153 a.a) and a putative Walker B motif (410–426 a.a) the amino acid sequences for which are shown in Table 5. These motif signatures seem to be more closely related to the metazoan/higher eukaryotic sequences.

**Table 5.**
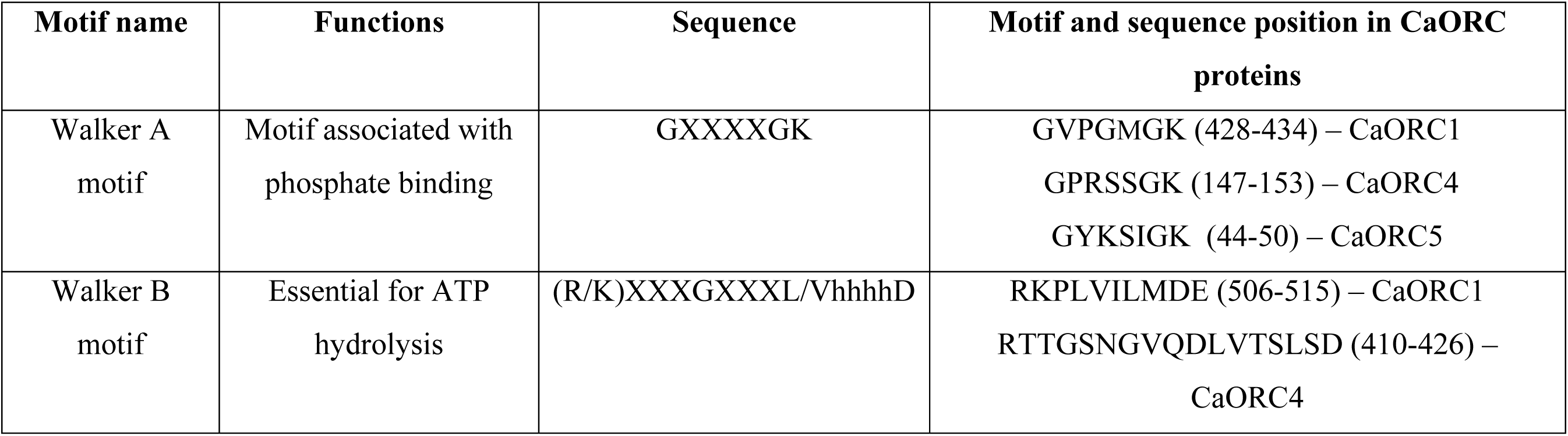
Putative signature of Walker motifs in CaORC proteins

### AT-hook motifs in CaORC proteins

AT hooks are DNA-binding motifs with a preference for A/T rich regions. These motifs are found in a variety of proteins, including the high mobility group (HMG) proteins. The AT-hook is a small motif which has a typical sequence pattern centered on a glycine-arginine-proline (GRP) tripeptide ^28,29^. The importance of this short conserved sequence is that it is necessary and sufficient for binding DNA and ori-ORC association. CaORC2 is found to have an AT-hook motif (182–197a.a, Figure 1D, Table 6) indicative of its propensity to bind origins.

**Table 6.** Domains of *C.albicans* ORC proteins compared with other eukaryotes

| Protein | *C.albicans* | *S.cerevisiae* | *S.pombe* | *D.melanogaster* | *X.laevis* | *M.musculus* | *H.sapiens* | *A.thaliana* |
| --- | --- | --- | --- | --- | --- | --- | --- | --- |
| ORC1 | BAH domain, PIP motif, AAA ATPase, Walker A & B motifs | BAH domain, AAA ATPase | BAH domain | BAH domain, AAA ATPase | BAH domain, AAA ATPase, PEST motif | BAH domain, AAA ATPase, PEST motif | BAH domain, AAA ATPase, PEST motif | BAH domain, PHD zinc finger, AAA ATPase, PEST motif |
| ORC2 | AT hook, PEST motif | AT hook | Not determined | No hits | No hits | No hits | No hits | PEST motif |
| ORC3 | MIR, PEST motif | ND | Not determined | AAA ATPase (P loop) | Not determined | No hits | MIR | Domain 1 Cullins, PEST motif |
| ORC4 | AAA ATPase, Walker A & B motifs | No hits | AT hook | AAA ATPase (P loop) | AAA ATPase (P loop) | AAA ATPase (P loop) | AAA ATPase (P loop) | AAA ATPase |
| ORC5 | WalkerA motif | AAA ATPase (P loop) | Not determined | AAA ATPase (P loop) | Not determined | AAA ATPase (P loop) | AAA ATPase (P loop) | AAA ATPase, PEST motif |
| ORC6 | No hits | No hits | Not determined | No hits | Not determined | No hits | No hits | No hits |
| CDC6 | AAA ATPase | AAA ATPase | Not determined | Not determined | AAA ATPase | AAA ATPase | AAA ATPase | AAA ATPase |
| CDT1 | No hits | Not determined | Not determined | No hits | No hits | No hits | No hits | PEST motif |

### PIP motif in CaORC proteins

A conserved Proliferating Cell Nuclear Antigen (PCNA) binding motif called the PCNA-interacting protein (PIP) box (QXXMXXFFFY) is found in the CaORC1 protein (524–536 a.a). Of the CaORC proteins, the PIP box is found to be unique to CaORC1.

### MOD1 motif in CaORC3

Two independent domains in human ORC3, a coiled-coil domain at the N terminus and a second region containing a MOD1-interacting region (MIR; 213–218aa)^30^, were found to be directly bound to the heterochromatin protein, HP1α^31^. A conserved peptide motif named MIR (MOD1 interacting region - PXVHH) which is essential for their interaction with MOD1, a serotonin-gated chloride channel that modulates locomotory behavior in *C. elegans^32^* is found in CaORC3 protein (435–448 a.a).

Although the CaORC proteins share less amino acid sequence homology with the ORC proteins of *S. cerevisiae* and *S. pombe*, the other ORC associated proteins (MCM proteins) seem to have higher homology (Table 7). Interestingly, the predicted molecular weights of the ORC complexes are equal in these three yeasts (Table 8).

**Table 7.**
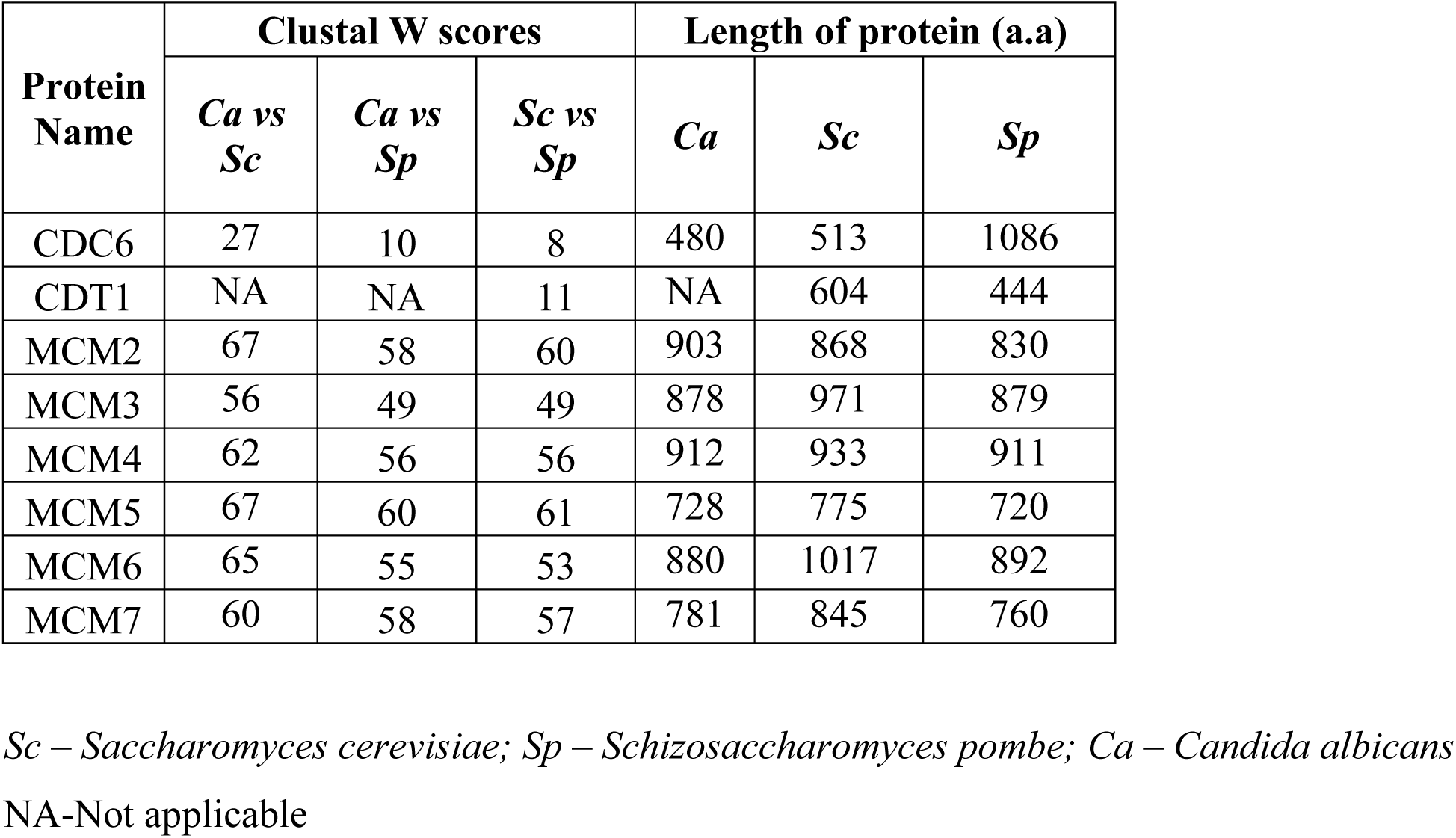
Comparison of the Clustal W scores and lengths of the ORC associated proteins in *S. cerevisiae* and *S. pombe* with *C. albicans*.

**Table 8.** Predicted molecular weight of the ORC proteins in *C. albicans, S. cerevisiae* and *S. pombe*

| ORC proteins | M.W in *S.cerevisiae*<br>(in KDa) | M.W in *S.pombe*<br>(in KDa) | M.W in *C.albicans*<br>(in KDa) |
| --- | --- | --- | --- |
| ORC1 | 120 | 80 | 91 |
| ORC2 | 72 | 61 | 78.6 |
| ORC3 | 62 | 80 | 79.2 |
| ORC4 | 56 | 108 | 64 |
| ORC5 | 53 | 52 | 57 |
| ORC6 | 50 | 31 | 41 |
| **Total** | **~412** | **~412** | **~412** |

### PEST motif in CaORC proteins

A PEST sequence is a peptide sequence that is rich in proline (P), glutamic acid (E), serine (S), and threonine (T). This sequence is associated with proteins that have a short intracellular half-life; hence, it is hypothesized that the PEST sequence acts as a signal peptide for protein degradation. CaORC2 (130–172 a.a.) and CaORC3 (6–33 a.a.) contain PEST motif. Analysis of PEST signals in human and mouse ORC proteins suggests that only ORC1 is targeted for ubiquitination which is likely to hold good for all mammals^33^. The domains of CaORC proteins are compared with other eukaryotes and are tabulated in Table 6 and compared with *S. cervevisiae* in Figure 1D. Recent studies have shown the evolution of the phospho regulation pattern in replication proteins of various yeast species including *C. albicans*^34^.

### Evolutionary relationships of ORC proteins

Molecular Evolutionary Genetics Analysis (MEGA) is an integrated tool for conducting sequence alignment, inferring phylogenetic trees, estimating divergence times, mining online databases, estimating rates of molecular evolution, inferring ancestral sequences, and testing evolutionary hypotheses^35^. The evolutionary history was inferred using the Neighbor-Joining method^36^. The optimal tree with the sum of branch length = 29.06450731 is shown in Figure 1A. The ORC1, ORC2 and ORC5 proteins from yeast to humans are found to have common nodes. Subsequently, the ORC proteins from the related species of *C. albicans* in the CTG clade were also compared and a phylogenetic tree was constructed (Figure 2A). The time tree demonstrates the diversification rate of these ORC proteins across the species of which ORC1 and ORC4 seem to be older than their counterparts (Figure 2B). In order to understand the sequence identity of the ORC sequences across various yeast species, Sequenceserver (http://blast.wei.wisc.edu/)^37,38^ was used across 86 publicly available yeast genomes (Figure 3; Supplementary Table 1).

**Figure. 2.**
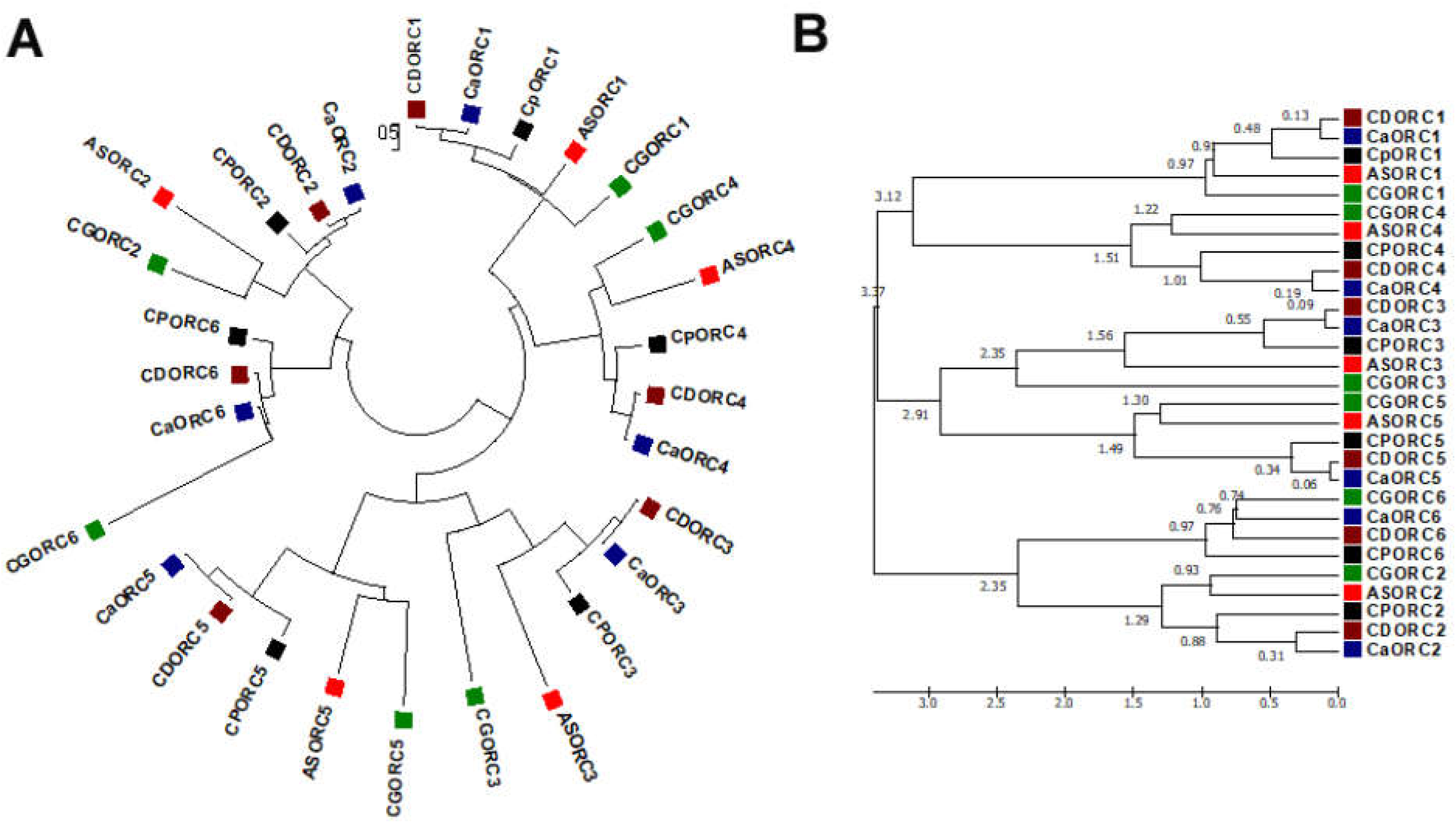
ORC phylogeny in CTG clade. **(A)** Molecular Phylogenetic analysis of ORC proteins in the CTG clade by Maximum Likelihood method. The evolutionary history was inferred by using the Maximum Likelihood method based on the JTT matrix-based model^75^. The tree with the highest log likelihood (−14518.9956) is shown. Initial tree**(s)** for the heuristic search were obtained automatically by applying Neighbor-Join and BioNJ algorithms to a matrix of pairwise distances estimated using a JTT model, and then selecting the topology with superior log likelihood value. The tree is drawn to scale, with branch lengths measured in the number of substitutions per site. The analysis involved 29 amino acid sequences. All positions containing gaps and missing data were eliminated. There were a total of 237 positions in the final dataset. Evolutionary analyses were conducted in MEGA6 ^80^. **(B)** The time tree molecular Phylogenetic analysis of ORC proteins in the CTG clade by the Maximum Likelihood method. The timetree shown was generated using the RealTime method ^81^. Divergence times for all branching points in the topology were calculated using the Maximum Likelihood method based on the JTT matrix-based model^79^. The estimated log likelihood value of the topology shown is −14518.9956. The tree is drawn to scale, with branch lengths measured in the relative number of substitutions per site. The analysis involved 29 amino acid sequences. All positions containing gaps and missing data were eliminated. There were a total of 237 positions in the final dataset. Evolutionary analyses were conducted in MEGA6^80^.

**Figure. 3.**
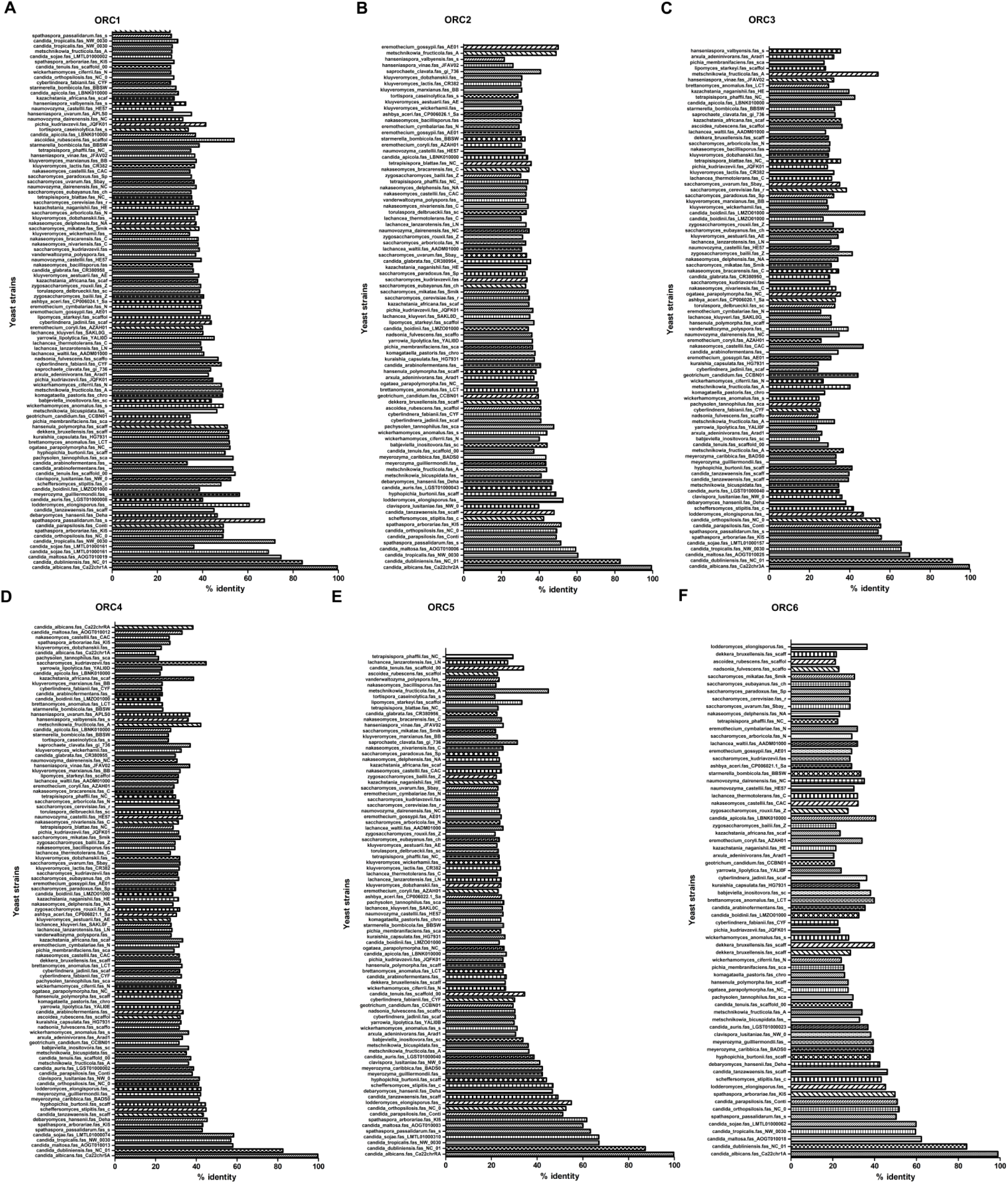
Comparative profile of percentage identity of CaORC proteins with other yeasts. The hexameric ORC complex containing ORC1-6 protein sequences are compared individually with diverse yeast species whose genome database is publicly available^37–38^ and their percent identity is plotted using Graphpad Prism^76^. **(A)** Percent identity of 104 hits of ORC1 sequences. **(B)** Percent identity of 86 hits of ORC2 sequences. **(C)** Percent identity of 90 hits of ORC3 sequences **(D)**. Percent identity of 104 hits of ORC4 sequences. **(E)**Percent identity of 88 hits of ORC5 sequences. **(F)** Percent identity of 69 hits of ORC6 sequences.

### Structure prediction of CaORC proteins

#### Prediction of secondary structure using Phyre

Over the past few decades, a number of computational tools for protein structure prediction have been developed. The **p**rotein **h**omology/analogy **r**ecognition **e**ngine (**Phyre**) is one of the widely used structure prediction systems providing a simple interface to results. The Phyre server (http://www.imperial.ac.uk/phyre) uses a library of known protein structures taken from the Structural Classification of Proteins (SCOP) database^39^ and augmented with newer depositions in the Protein Data Bank (PDB)^40^. The sequence of each of these structures is scanned against a non-redundant sequence database and a profile is generated and deposited in the ‘fold library’. The known and predicted secondary structure of these proteins is also stored in the fold library. The popular web servers for fold recognition are Phyre, I-TASSER, SAM-T06, HHpred.

We used I-TASSER (Iterative Threading ASSEmbly Refinement^41^) for structure prediction of CaORC proteins. Of all the CaORC proteins, CaORC4 was found to be one of the putative candidates for further fine refinement studies of the protein structure due to its higher Cscore (combined measure, See Methods section) which indicates a better confidence in predicting the function using the template (Table 9). Hence, we proceeded for predicting the structure of CaORC4 using Phyre.

**Table 9.** Cscore values of CaORC proteins

| **ORC proteins** | **Cscore<sup>GO</sup>** | **Cscore<sup>LB</sup>** |
| --- | --- | --- |
| ORC1 | 0.24 | 0.41 |
| ORC2 | 0.25 | 0.02 |
| ORC3 | 0.16 | 0.01 |
| ORC4 | 0.29 | 0.6 |
| ORC5 | 0.29 | 0.58 |
| ORC6 | 0.21 | 0.01 |

### Secondary structure and disorder prediction for CaORC4

The query sequence (CaORC4p) is scanned against the non-redundant sequence database and a profile is constructed. Five iterations of PSI-BLAST are used to gather both close and remote sequence homologs. The PSI-BLAST provides a means of detecting distance relationships between proteins. The pair-wise alignments generated by PSI-BLAST are combined into a single alignment with the query sequence as the master. The secondary structure of CaORC4p is predicted following profile construction.

Three independent secondary structure prediction programs are used in Phyre: Psi-Pred1^42^, SSPro^43^ and JNet^44^. The output of each program is in the form of a three-state prediction: alpha helix, beta strand and coil. Each of these three programs provides a confidence value at each position of the query for each of the three secondary structure states. These confidence values are averaged and a final, consensus prediction is calculated and displayed beneath the individual predictions.

### Fold recognition for CaORC4

The profile and secondary structure of CaORC4 is then scanned against the fold library using a profile-profile alignment algorithm detailed in^45^. This alignment process returns a score on which the alignments are ranked. These scores are fitted to an extreme value distribution to generate an E-value. The top ten highest scoring alignments are then used to construct full 3D models of the CaORC4p (Figure 4A and Figure 4B).

**Figure. 4.**
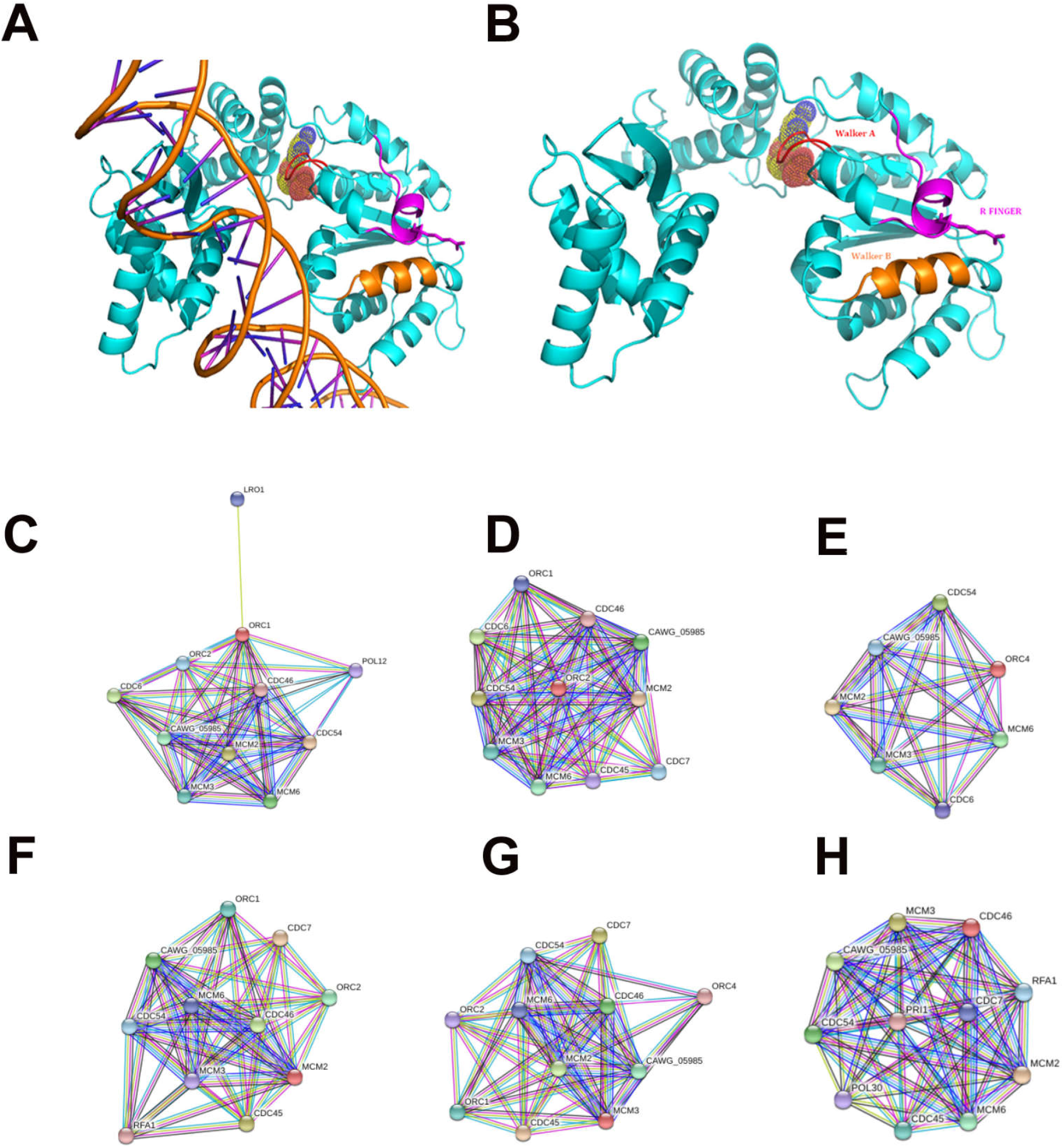
3D model of CaORC4 and putative interactors of CaORC proteins. **(A)** 3D model of CaORC4 with DNA. **(B)** 3D model of CaORC4 with Walker A bound to ATP sphere, Walker B and R finger motifs. (C-H) Protein interaction map of the *C. albicans* pre-RC including CaORC1, CaORC2, CaORC4, CaMCM2, CaMCM3, CaMCM5 (CDC46) respectively (Table 10). The bright red circle is the query protein. The interaction map of CaMCM4 and CaMCM6 are the same as CaMCM3.

### Interactions of pre-RC proteins - SMART prediction

SMART (<u>S</u>imple <u>M</u>odular <u>A</u>rchitecture <u>R</u>esearch <u>T</u>ool) is a web-based tool (http://smart.embl.de/) that allows rapid identification and annotation of protein domains and the analysis of protein domain architectures. This provides the complete set of protein descriptions allowing users to quickly find relevant information^46–47^. The predicted functional partners of the preRC proteins in *C. albicans* are enlisted in the Table 10 and are also shown schematically in the Figure 4C-H. In short, it is evident that although the size of the individual proteins in the ORC complex across diverse yeast species is varied, the whole complex constitutes to ~412 KDa (Table 8). Our *in silico* analysis suggests that although CaORC proteins share less sequence homology with yeasts, Drosophila, Xenopus, mouse and humans (Table 7), some of the characteristic functional motifs are retained in them (Figure 1D, Table 6). CaORC1 is found to have the BAH domain and the PIP motif, CaORC2 has an AT-hook motif, and CaORC3 has a MOD1-interacting region (MIR). The AAA ATPase is found in CaORC1 and CaORC4 and the PEST motif is found in CaORC2 and CaORC3. We used Phyre to predict the secondary structure and modeled the 3D structure of CaORC4 with walker A and B motifs and arginine finger motif. We used SMART predictions to check the putative interactive partners of CaORC proteins of which CaORC4 was found to have no direct interaction with any other CaORC protein.

**Table 10.** SMART predictions of pre-RC proteins’ interactions

| **Pre-RC protein** | **Domains / motifs** | **Putative interacting partners** |
| --- | --- | --- |
| CAORC1 | BAH & AAA | ORC2,CDC6,CDC46,<u>CDC54,MCM2,MCM3,MCM6, CAWG05_985</u>,POL12,LRO1 |
| CAORC2 |  | ORC1,CDC6,CDC46,<u>CDC54</u>,CDC45,CDC7,<u>MCM2, MCM3,MCM6,CAWG05_985</u> |
| CAORC3 |  | - |
| CAORC4 | AAA | <u>CDC54</u>,CDC6,<u>MCM2,MCM3,MCM6,CAWG05_985</u> |
| CAORC5 |  | - |
| CAORC6 |  | - |
| CACDC6 | AAA | - |
| CAMCM2 | MCM | ORC1,ORC2,CDC45,CDC46,<u>CDC54</u>,CDC7,<u>MCM3, MCM6,CAWG05_985</u>,RFA1 |
| CAMCM3 | AAA / MCM | ORC1,ORC2,ORC4,CDC45,CDC46,<u>CDC54</u>,CDC7,<u>MCM2, MCM3,MCM6, CAWG05_985</u> |
| CAMCM4 / CDC54 | MCM | ORC1,ORC2,CDC45,CDC46,<u>CDC54</u>,CDC7,<u>MCM2, MCM3,MCM6, CAWG05_985</u> |
| CAMCM5 / CDC46 | AAA / MCM | CDC45,<u>CDC54</u>,CDC7,<u>MCM2,MCM3,MCM6, CAWG05_985</u>,PRI1, POL30, RFA1 |
| CAMCM6 | MCM | ORC1,ORC2,ORC4,CDC45,CDC46,**<u>CDC54</u>**,CDC7,**<u>MCM2</u>**,<br>**<u>MCM3,MCM6, CAWG05_985</u>** |
| CAMCM7 | AAA /<br>MCM | - |

Note : CAWG05\_985 = MCM7

## Discussion

CaORC proteins (CaORC1-6) and their associated proteins were identified by a BLAST analysis using the *S. cerevisiae* proteins as the query sequences in the Candida Genome Database (CGD). The phylogenetic analysis suggests that in spite of limited amino acid sequence similarity with their counterparts in other organisms, the CaORC proteins share most of the functional domains with them. Interestingly, the amino acid sequences of CaORC1, 2 and 4 share higher degree of similarities than CaORC3, 5 and 6 to those of *S. cerevisiae*. CaORC1, 4 and 5 tend to be homologous to the mammalian counterparts. Moreover, the CaORC proteins are also compared across CTG clade and other yeast species to provide a robust roadmap for further comparative yeast subphylum analysis (Figure 2 and Figure 3). Of the other preRC components compared here, Cdt1 has no apparent homolog in *C. albicans*, whereas, all other pre-RC proteins such as Cdc6 and Mcm2-7are very similar to their counterparts in other yeasts.

The main function of ORC proteins is to associate specifically with origins and recruit other factors including Cdc6 and MCM2-7 to form the preRC. In *S. cerevisiae*, the origins have a conserved stretch of 11 bp, the ARS consensus sequence, ACS, which is essential for ORC binding and origin activity. The replication origins of *C. albicans* (based on limited available data^48–51^) appear to be similar to those of *S. pombe* and other higher eukaryotes in having no such consensus sequence. In *S. pombe*, ORC4 binds with AT-rich origins via its 9 AT-hook motifs. Moreover, the ORC-origin binding might be affected both by intrinsic factors such as the DNA sequence that marks the ORC binding site and by extrinsic factors such as the chromatin component that marks both the histone and non-histone proteins. The absence of conserved sequences in *C. albicans* origins^50^ along with our *in silico* analysis suggests that the CaORC-origin interactions would be largely chromatin dependent. In the genome-wide studies for identification of replication origins in *C. albicans* by ChIP-microarray based approach using an antibody against *S. cerevisiae* ORC complex, low nucleosome occupancy has been shown as conserved landmark of replication origins in *C. albicans*^51^.

The BAH module found in several chromatin-associated proteins play important roles in gene silencing, replication and transcriptional regulation by promoting protein-protein interaction^12^. The BAH domain of humanORC1 has been shown to bind to H4K20me2^52^ and abrogation of this binding causes impaired ORC1 loading onto origins, and cell cycle progression. The BAH domain present in CaORC1 along with the highly conserved basic residues (K-362 and R-367)^53^ in its AAA domain is likely to play a key role in ORC-origin binding in *C. albicans*.

The AAA+ domains present in different ORC subunits (ORC1 and 5 in *S. cerevisiae* and ORC1, 4, and 5 in metazoans) are important for the assembly of ORC at origins and those in Cdc6 are critical for the loading of the MCM proteins (Table 6). Like metazoans, the CaORC subunits 1, 4, and 5 and CaCdc6 containing AAA+ domains are likely to be engaged in ORC assembly and consequent MCM recruitment although a perfect match to the Walker A and B motifs are present only in CaORC4 (Table 4 and 5). In all tested organisms, ORC1 has been found to be the major ATPase required for ORC assembly at origins. Experimental evidence would be required to find out if some or all of these subunits are involved in ATP binding and hydrolysis in *C. albicans*. Cdt1 helps in Cdc6 recruitment to origin bound ORC and is important in cell cycle regulation of preRC assembly at origins and limiting replication to a single round per cell cycle. The absence of a Cdt1 homolog in *C. albicans* suggests that this important task may be accomplished by a different mechanism/factor (Table 6 and Table 7). The unique presence of the PEST motif in CaORC2 and CaORC3 indicates that these components might be degraded in a cell cycle specific manner facilitating ORC turnover. The unique sequence of nine copies of AT hook motifs present in SpORC4 are critical for origin binding of ORC4 which is ATP-independent in *S. pombe*^24^. In *S. cerevisiae*, the origin binding of ORC is ATP-dependent and the presence of single DNA-binding AT-hook motif (PRKRGRPRK) is identified in the disordered regions of ScORC2^19^. The presence of the small AT-hook motif in CaORC2 to be another plausible motif for origin binding and their role in replication remains highly speculative. Similarly, it remains elusive as to whether the presence of MIR domain in CaORC3 has any role in silencing by binding to any heterochromatin component like HP1.

In *S. cerevisiae*, the ScORC proteins associate with origins in a sequence-dependent manner. Only ORC1, ORC2, ORC4 and ORC5 appear to make direct contacts to A and B1 domains of the replication origin^33, 54–55^. ScORC3 helps in forming the stable complex without directly binding to the DNA whereas ORC6 does not bind to the DNA but helps in recruiting multiple Cdt1 molecules^56–59^. The situation is very different in *Drosophila* cells where DNA replication initiates at many sites, which are probably sequence independent, throughout the genome at the same time^60^. In contrast to ScORC6, which is not required for DNA binding, DmORC6 is required for the DNA binding of DmORC and is an integral part of the DmORC complex^61^. The DmORC6 alone has DNA binding activity, likely due to the predicted TFIIB-like DNA binding domain in the smallest subunit^62^. DmORC binds DNA with little sequence specificity. ORC proteins generally require ATP to interact specifically with origin DNA (except in the case of SpORC). In all the species studied so far, ORC1, ORC4 and ORC5 contain potential ATP binding sites. ATP hydrolysis by ORCs to regulate DNA binding is well studied in ScORCs and DmORCs^26, 59^. The *in silico* predictions by Beltrao and colleagues^34^ showed the increasing probability of CaORC2, CaORC4 and CaORC6 proteins to be phosphorylated by Cdc28, a cyclin dependent protein kinase.

We were able to build the 3D protein structure of CaORC4 only whereas the other CaORC proteins did not have good homology with the known PDB (Protein Data Bank) structures. From our *in silico* analysis of interactive studies, it is evident that CaORC3, CaORC5 and CaORC6 do not interact with the other ORC counterparts. It is possible that only CaORC1, CaORC2 and CaORC4 would be involved in DNA binding during the process of DNA replication and the other counterparts may aid in tethering or in conformational organization. CaCdc6 and Cdc54, the apparent common binding partners of CaORC1, CaORC2 and CaORC4 and many MCMs are also predicted to play important role(s) in preRC assembly and functioning. We also find a potential ATP binding site in CaORC4 which might help in the regulation of origin binding. The mode of ORC assembly at origins in *C. albicans* might be different from that in other yeasts. The *in silico* detection of the presence of AAA+ ATPase and Walker motifs in CaORC4 and its likely interaction with MCM proteins suggest that CaORC4 might be involved in stable binding to origin DNA and loading MCM proteins to origins. While possibilities of a physical association between CaORC4 and other CaORC proteins were not obvious, the role of some unknown factors mediating ORC assembly in *C. albicans* is not ruled out. CDC6, CDC54 and MCM proteins interact with CaORC1, CaORC2 and CaORC4. In absence of a direct interaction of CaORC4 with other ORC counterparts, these proteins might be mediating interaction between them. Moreover, the absence of Cdt1 in *C. albicans* might provide an additional role for CaORC4.

Recent studies demonstrate that besides the involvement of specific proteins that control DNA replication, some enzymes with primary functions that are involved in various other processes can also play a vital role in the regulation of genome duplication. There seems to be a direct link between central carbon metabolism and DNA replication regulation from prokaryotes ^63–65^ to eukaryotes including humans^66–67^. A recent analysis^66^ demonstrates that partial silencing of genes encoding for the glycolytic and TCA enzymes affects the entry of human fibroblasts into the S-phase. It is also reported that ScORC proteins interact with some of the metabolic genes that are associated with replication origins^68^. One such example is the hexokinase (HXK2) gene which at a decreased level causes substantial impairment in DNA synthesis. Our preliminary reports from Co-IP studies (data not shown) also showed an interaction of CaORC4 with HXK2 by which it is speculated that CaORC4 might play a role in the regulation of central carbon metabolism besides its cardinal role of DNA replication. This can be further supported by the induced expression of CaORC4 in response to alpha pheromone in SpiderM medium^69^.

From the above observations, we hypothesize that CaORC4 might be less tightly associated with the core preRC complex but involved in cell cycle regulation and DNA checkpoint activation. It is quite possible that CaORC4 may not be bound to chromatin throughout the cell cycle as seen in *Drosophila* and yeast ^33^. Recent studies advocate a concerted interaction between ORCs, nucleosomes and replication origin DNA that stabilizes ORC-origin binding in yeast. The atomic force microscope (AFM) studies show that ORC establishes its origin interaction by binding to both nucleosome-free origin DNA and neighboring nucleosomes that are species-specific^70^.

Recent reports suggest that Drg1, an AAA-ATPase protein is the potential target for the drug diazaborine. This drug is demonstrated to block ribosome biogenesis in yeast^71^. Similarly, a valosin containing AAA-ATPase protein, P97 is found to be a therapeutic target for CB-5083 in the cancer treatment^72^. A study on Trypanosoma ORC has raised possibilities on identifying novel drug targets demonstrating the drug potential of the pre-replication machinery^73^.

Our *in silico* studies would form the basis for understanding the domain architecture and further characterization of CaORC proteins which can be validated by *in vitro* studies. It may ultimately provide clues about the potential drug targets helping curb Candida infection at the step of DNA replication.

## Methods

### Annotation of *C. albicans* pre-RC genes

The genome of *C. albicans* (http://www.candidagenome.org/) was searched for homologs of pre-RC complex genes using BLAST^74^. Alignment of pre-RC gene sequences from Candida and its homologs in other eukaryotic organisms was carried out using the ClustalW algorithm^15^.

The pairwise ClustalW scores are calculated by the number of identities between the two sequences, divided by the alignment length in terms of percentage.

### Phylogenetic analysis

Phylogenetic analysis was performed with the MEGA4 program^75^.

### *In silico* analysis

The putative protein sequences whose theoretical characteristics were obtained using several programs in the ExPASy (Expert Protein Analysis System) server of the Swiss Institute of Bioinformatics (www.expasy.ch/tools/). Protein sequences were entered into MotifScan (pattern searches), ProDOM (protein domain identification), Interpro (protein domain and pattern search identification), NetPhos (prediction sites for phosphorylation) and PESTfind (identification of PEST sequences), SMART (prediction of protein domain architecture) and Phyre (secondary structure prediction). To determine the sequence identity of CaORC across 86 diverse publicly available yeast databases, a TBLASTN was performed in the Sequenceserver (http://blast.wei.wisc.edu/) with CaORC proteins as the query sequence^37–38^ and the percent identity was plotted against the species using Graphpad Prism^76^.

### Phyre structure prediction parameters

Cscore^GO^ is a combined measure for evaluating global and local similarity between query and template protein. This score ranges from 0–1 where a higher value indicates a better confidence in predicting the function using the template. Cscore^LB^ is the confidence score of predicted binding site of the protein with values ranging between 0–1. Higher the score more reliable is the ligand binding prediction.

## Acknowledgements

This work was supported by Department of Biotechnology to KS and DDD (BT/PR13724/BRB/10/782/2010). The award of direct Senior Research Fellowships to SP from Council of Scientific and Industrial Research (9/1014(0001)2K10-EMR-I) is greatly acknowledged. We thank Dr. E.J.Woo, Korea for the 3D structural studies.

## Author Contributions

S.P. performed experiments, analyzed data and wrote the paper. K.S. and D.D. designed the study, analyzed data, and wrote the paper.

